# Tracking Inflammation and Fibroblast Activation in Hypertensive Heart Failure Across the Cardio-Renal Axis

**DOI:** 10.64898/2026.07.13.738150

**Authors:** Maja Strunk, Annika Hess, Marcel Gutberlet, Michael Willmann, Tobias L Ross, Frank M Bengel, James T Thackeray

## Abstract

Hypertension and heart failure are associated with increased risk of chronic kidney disease. Cardiorenal syndrome is characterized by excessive systemic inflammation and progressive fibrosis. We hypothesized that transient hypertension in mice due to infusion of angiotensin II and phenylephrine (Ang/Phe) would induce parallel immune cell and fibroblast activation in both the heart and kidney, where the intensity of inflammation and fibroblast activity would predict decline in function of both organs. Adult male C57Bl/6N mice were randomized to receive 7d infusion of either Ang/Phe (n=41) or vehicle (n=27) by subcutaneous osmotic minipump. Despite removal of minipumps at 7d, Ang/Phe mice displayed persistent myocyte hypertrophy, interstitial fibrosis, and modestly reduced systolic function to 6 weeks. Molecular imaging of chemokine receptor CXCR4 using 68Ga-pentixafor at 3d of Ang/Phe infusion revealed transient inflammation in the left ventricle. Imaging of fibroblast activation protein (FAP) revealed diffuse fibroblast activity in the left ventricle. Both imaging signals predicted subsequent functional decline. Magnetic resonance imaging of the kidney revealed transient prolongation of T1 relaxation in at 2 weeks after Ang/Phe infusion that returned to normal by 6wk, despite a progressive reduction in renal perfusion. CXCR4 and FAP PET displayed no change in kidney inflammation or fibroblast activation. Comparison of imaging data described a direct correlation between cardiac and renal CXCR4 PET signal at 3d and FAP PET signal at 7d. The intensity of cardiac inflammation correlated with subchronic fibrosis in the kidney cortex. Total body molecular imaging enables simultaneous evaluation of the immune-fibrosis network in heart-kidney crosstalk after short term hypertension and may provide valuable guidance of novel therapies.

## INTRODUCTION

Cardiovascular disease and heart failure remain the leading cause of global mortality and represent a major burden to health care systems worldwide ^1^. The consequences of cardiac disease are not restricted to the heart. In particular, hypertension and heart failure are associated with higher incidence of chronic kidney disease ^2,3^, but the bidirectional pathobiology of the heart-kidney axis remains incompletely understood.

Cardio-renal syndrome can develop with primary injury to the heart or the kidney, and imparts heightened risk of failure to the reciprocal organ. Declining glomerular filtration rate independently predicts re-hospitalization for congestive heart failure ^2,4^. Moreover, up to 40% of patients with acute MI and 50% of patients with heart failure with preserved ejection fraction exhibit chronic kidney disease ^5–7^. Both heart failure and kidney disease impart a systemic pro-inflammatory state, which may influence other organ systems ^3^.

Medical imaging can provide insights into the pathogenic mechanisms linking the heart and kidney. Reduced renal perfusion assessed by ^13^N-ammonia positron emission tomography is associated with added risk of cardiovascular events and renal outcomes ^8^. In rodent models of myocardial infarction, increased inflammation in the heart is paralleled by a concurrent increase in inflammation avid radiotracer retention in the kidney ^9^.

Moreover, organ fibrosis is central to the progression of cardio-renal syndrome ^10^. Single cell sequencing studies have identified commonalities in heart and kidney failure specifically relating to markers of organ fibrosis ^11–13^. Notably, the expression of fibroblast activation protein (FAP) by transiently activated cardiac fibroblasts has been reported in persistent hypertension ^14^, and are intrinsically linked to local immune cell activity ^15,16^.

Given the synergy in disease progression, we hypothesized that persistent hypertension would precipitate sequential inflammation and fibroblast activation in both the heart and kidney. We induced hypertensive heart failure by temporary subcutaneous infusion of angiotensin II and phenylephrine in mice and measured acute inflammation and subacute fibroblast activation using total body PET imaging of chemokine receptor CXCR4 and FAP, respectively. Cardiac and renal tissue characterization was validated by magnetic resonance imaging and histopathology.

## RESULTS

### Transient Ang/Phe infusion generates persistent left ventricle remodeling

To induce hypertensive heart failure, mice received continuous Ang/Phe infusion via subcutaneous osmotic minipump over 7d. Left ventricle mass was elevated in the first days of infusion, with a 40% increase in necropsy mass evident at 3d which was further exacerbated by 7d (Fig 1A). The Ang/Phe stimulus was removed from 7d, but ventricle hypertrophy continued to increase over 6 weeks, as evidenced by serial CMR ventricle mass measurements (Fig 1B). WGA staining confirmed elevated cardiomyocyte cross-sectional area from 3d of Ang/Phe infusion which remained to 6 weeks (Fig 1C). This ventricle remodeling resulted in an increase in end systolic and end diastolic volume, first evident from 2 weeks and growing over 6 weeks (Fig S2). CMR revealed modestly impaired contractile function (Fig 1D), with a gradual decline in left ventricle ejection fraction (Fig 1E). Picrosirius red staining identified diffuse interstitial fibrosis throughout the left ventricle from 3d of Ang/Phe infusion, reaching a maximum at 7d (Fig 1G). Removal of the Ang/Phe pump alone did not reverse fibrosis (Fig 1EF). The extent of fibrosis at 6 weeks inversely correlated with percent fibrotic area on necropsy (Fig 1H).

**Figure 1.**
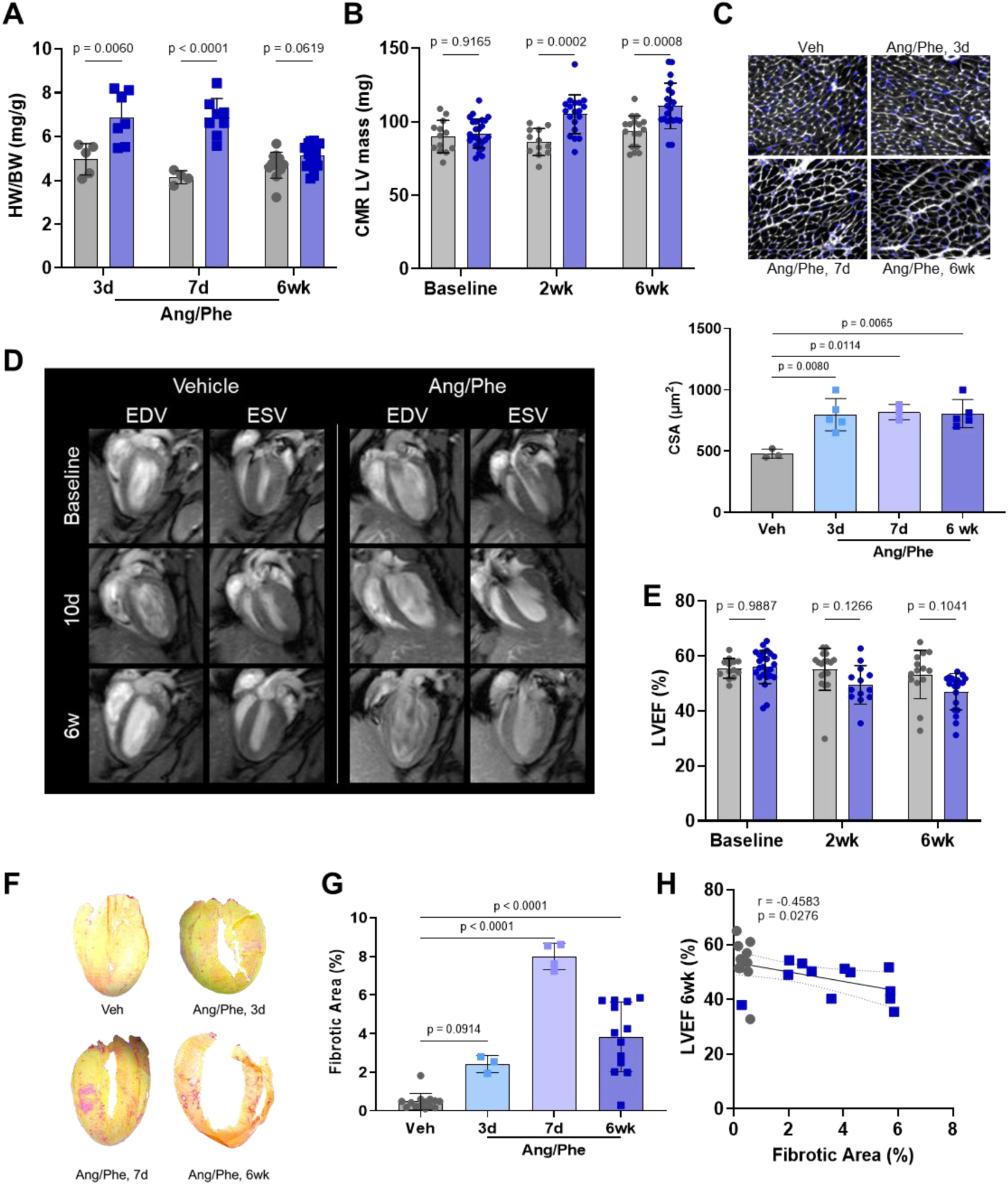
Cardiac function after transient Angiotensin/Phnylephrine infusion. (**A**) Heart weight to body weight (HW/BW) ratio on necropsy and (**B**) cardiac magnetic resonance (CMR) imaging estimation of left ventricle (LV) mass at intermediate intervals after 7d of Ang/Phe administration. (**C**) Wheat germ agglutinin staining identifies gradual increase in cardiomycyte cross-sectional area (CSA) after Ang/Phe. (**D**) Serial CMR images display end diastolic volume (EDV) and end systolic volume (ESV) with (**E**) moderate reduction in left ventricle ejection fraction (LVEF). (**F**) Picrosirius red staining and (**G**) quantification of fibrotic area after Ang/Phe. (**H**) Pearson correlation between extent of picrosirius red-defined interstitial fibrosis and LVEF. Statistics: mixed effects model for multiple comparisons in serial data; ordinary one-way ANOVA for biologic replicates in histology and necropsy; Pearson product-moment correlation for fibrotic area and LVEF.

### Cardiac inflammation and fibroblast activation after Ang/Phe infusion

To evaluate inflammation in the early stages of hypertension, animals underwent 68Ga-pentixafor imaging at 3d of Ang/Phe infusion. PET/CT images demonstrated global diffuse CXCR4 signal in the left ventricle compared to vehicle controls (Fig 2A), with a 50% increase in tracer accumulation (Fig 2B). Ex vivo autoradiography confirmed the signal localization to the myocardium (Fig S3) which was associated with increased density of CD68+ macrophages on immunostaining (Fig 2C).

**Figure 2.**
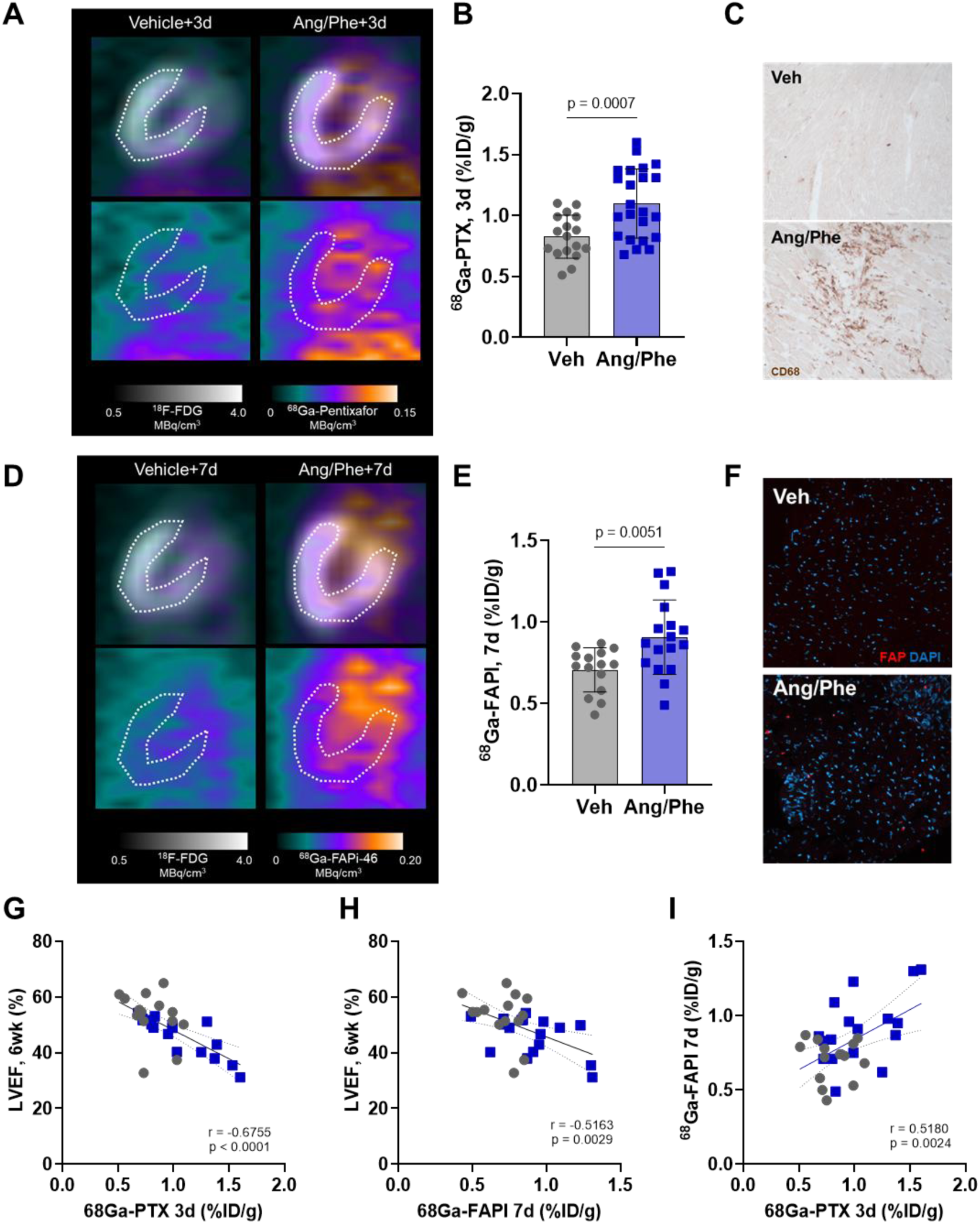
Molecular imaging of inflammation and fibroblast activation in the heart. (**A**) Representative images and (**B**) semi-quantification of ^68^Ga-pentixafor (PTX) uptake as % injected dose (ID)/g tissue in the left ventricle at 3d of angiotensin II/phenylephrine (Ang/Phe) infusion compared to vehicle (Veh) showing chemokine receptor CXCR4 signal (colourscale). Left ventricle contours are defined by subsequent viability imaging with ^18^F-fluorodeoxyglcuose (FDG, greyscale). (**C**) Immunostaining of CD68 at 3d. (**D**) Representative images and (**E**) semi-quantification of 68Ga-FAPI-46 (FAPI) uptake in the left ventricle at 7d of Ang/Phe or vehicle showing fibroblast activation protein (FAP, colourscale). Left ventricle contours are defined by subsequent viability imaging with ^18^F-FDG (greyscale). (**F**) Immunostaining of FAP validates tracer signal at 7d. Pearson correlation between (**G**) PTX signal at 3d and left ventricle ejection fraction (LVEF) at 6 wks, (**H**) FAPI signal at 7d and LVEF at 6 wks, and (**I**) PTX signal at 3d and FAPI signal at 7d. Statistics: unpaired t test with Welch correction for comparison of two groups, Pearson product-moment correlation between continuous variables.

Because hypertensive heart failure is associated with interstitial fibrosis, we next evaluated fibroblast activation protein by PET/CT imaging with 68Ga-FAPI-46 at 7d of Ang/Phe infusion. Ang/Phe infused mice showed a modest but significant increase in cardiac FAP signal, diffusely localized throughout the left ventricle (Fig 2D,E). Autoradiography confirmed the selective accumulation of ^68^Ga-FAPI-46 within the myocardium (Fig S3) which corresponded to regional increases in FAP immunostaining (Fig 2F). After removal of the Ang/Phe stimulus at 8d, we evaluated the persistence of FAP signal at 6 weeks of heart failure. There was no difference in quantitative FAP signal in the left ventricle at the chronic timepoint (Fig S4). Repeated imaging demonstrated that cardiac FAP signal declined from 7d to 6 weeks (Fig S4).

CXCR4 PET imaging signal at 3d directly correlated with left ventricle volumes at the chronic endpoint (Fig S5), and inversely correlated with chronic left ventricle ejection fraction (r=-0.6755, p<0.0001, Fig 2F). Similarly, the intensity of FAP PET signal at 7d predicted left ventricle geometry (Fig S5) and ejection fraction at 6 weeks (r=, -0.5163, p=0.0029, Fig 2G). Moreover, the intensity of inflammation at 3d correlated with FAP signal at 7d (r=0.5180, p=0.0024).

### Transient Ang/Phe infusion and heart failure leads to modest renal remodeling

Given the connection between hypertension, heart failure, and kidney function, we evaluated whether short-term Ang/Phe infusion influenced chronic kidney morphology. Necropsy examination demonstrated limited differences in kidney mass over the acute phase of disease (Fig 3A,B), which was attenuated when normalized to body mass (Fig S). We evaluated morphology changes in kidney by serial MR imaging prior to Ang/Phe, at 10d (3d after 7d infusion and pump removal) and at 6 weeks. T1 images revealed a transient prolongation of relaxation time in the subchronic phase in the renal cortex (Fig 3C,D) and global medulla (Fig 3E), which returned to baseline levels by 6 weeks. Similarly, T2 relaxation time was mildly prolonged at 2 weeks after Ang/Phe in both the renal cortex (Fig 3F,G) and medulla (Fig 3H). There was no temporal change in T1 or T2 relaxation time over the course of the experiment in vehicle-treated animals (Fig S6). Arterial spin labelling demonstrated a gradual decline in renal perfusion, reaching 80% of baseline at 6 weeks (Fig 3I,J). T1 relaxation time in the renal cortex directly correlated with renal perfusion on arterial spin labeling (Fig 3K).

**Figure 3.**
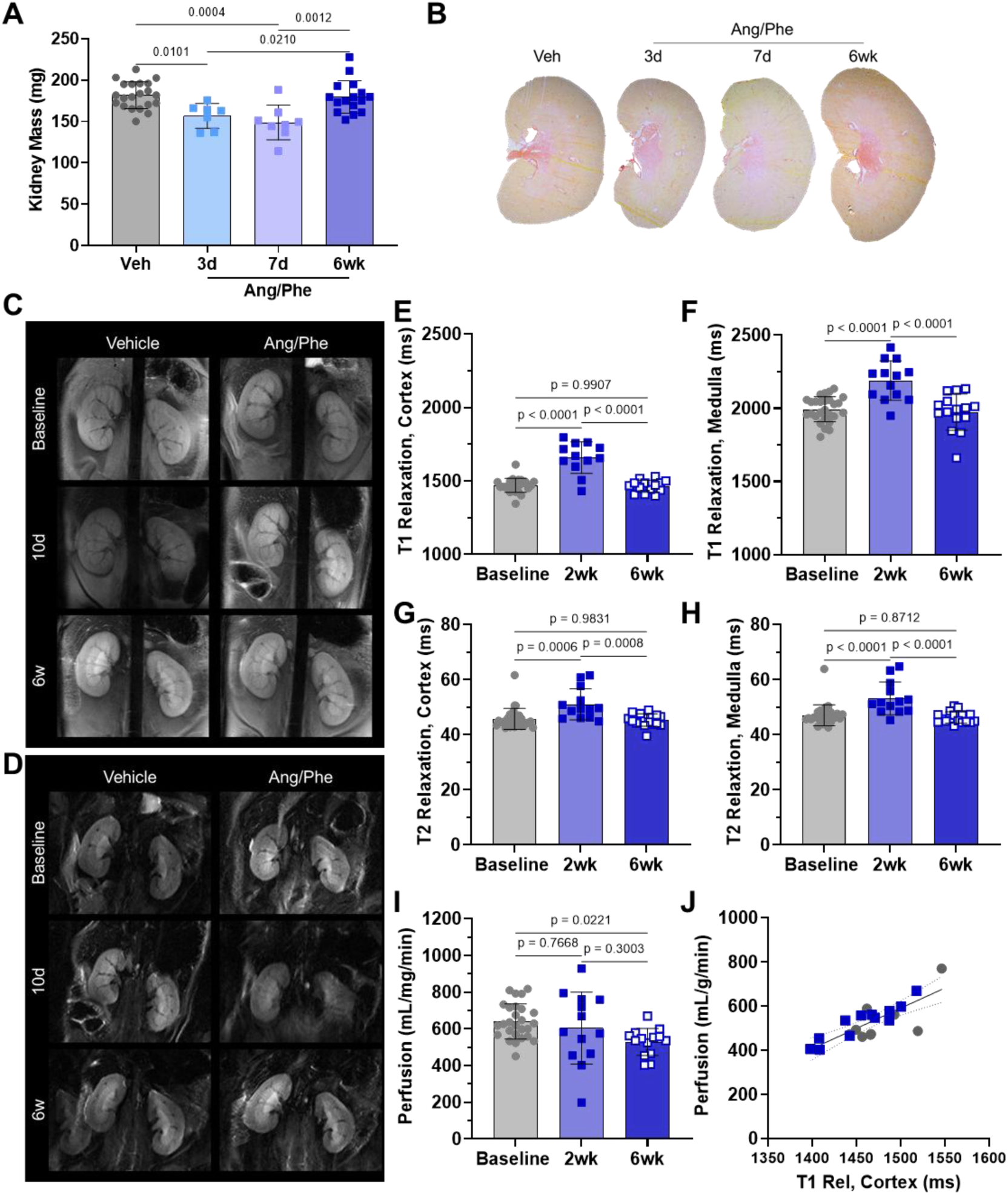
Renal function after transient Angiotensin/Phnylephrine infusion. (**A**) Necropsy measurements of kidney mass and (**B**) picrosirius red staining of fibrosis at intermediate timepoints of Ang/Phe or vehicle (veh) administration. (**C**) Serial T1-weighted magnetic resonance images of kidneys after Ang/Phe or Veh treatment and calculated relaxation (rel) time in (**D**) renal cortex and (**E**) renal medulla. (**F**) Serial T2-weighted magnetic resonance images of kidneys after Ang/Phe or Veh treatment and calculated relaxation time in (**G**) renal cortex and (**H**) renal medulla. (**I**) Arterial spin labeling derived renal perfusion after Ang/Phe or vehicle treatment. (**J**) Pearson correlation between T1 relaxation at 2wk and perfusion at 6wk in the renal cortex. Statistics: Ordinary one-way ANOVA for single timepoint histology comparison and serial magnetic resonance imaging measurements; Pearson product-moment correlation for comparison of continuous variables.

### Renal inflammation and fibroblast activation after Ang/Phe infusion

We then evaluated inflammation and fibroblast activation in the kidney by sequential PET imaging during Ang/Phe infusion. ^68^Ga-Pentixafor PET revealed no difference in kidney CXCR4 signal at 3d after Ang/Phe infusion (Fig 4A,B). Regional distribution was validated by ex vivo autoradiorgraphy, with higher tracer signal in the medulla, but no significant difference in tracer signal between Ang/Phe and vehicle controls (Fig S7). Immunohistochemistry revealed similar CD68+ cell content in the renal cortex (Fig 4C).

**Figure 4.**
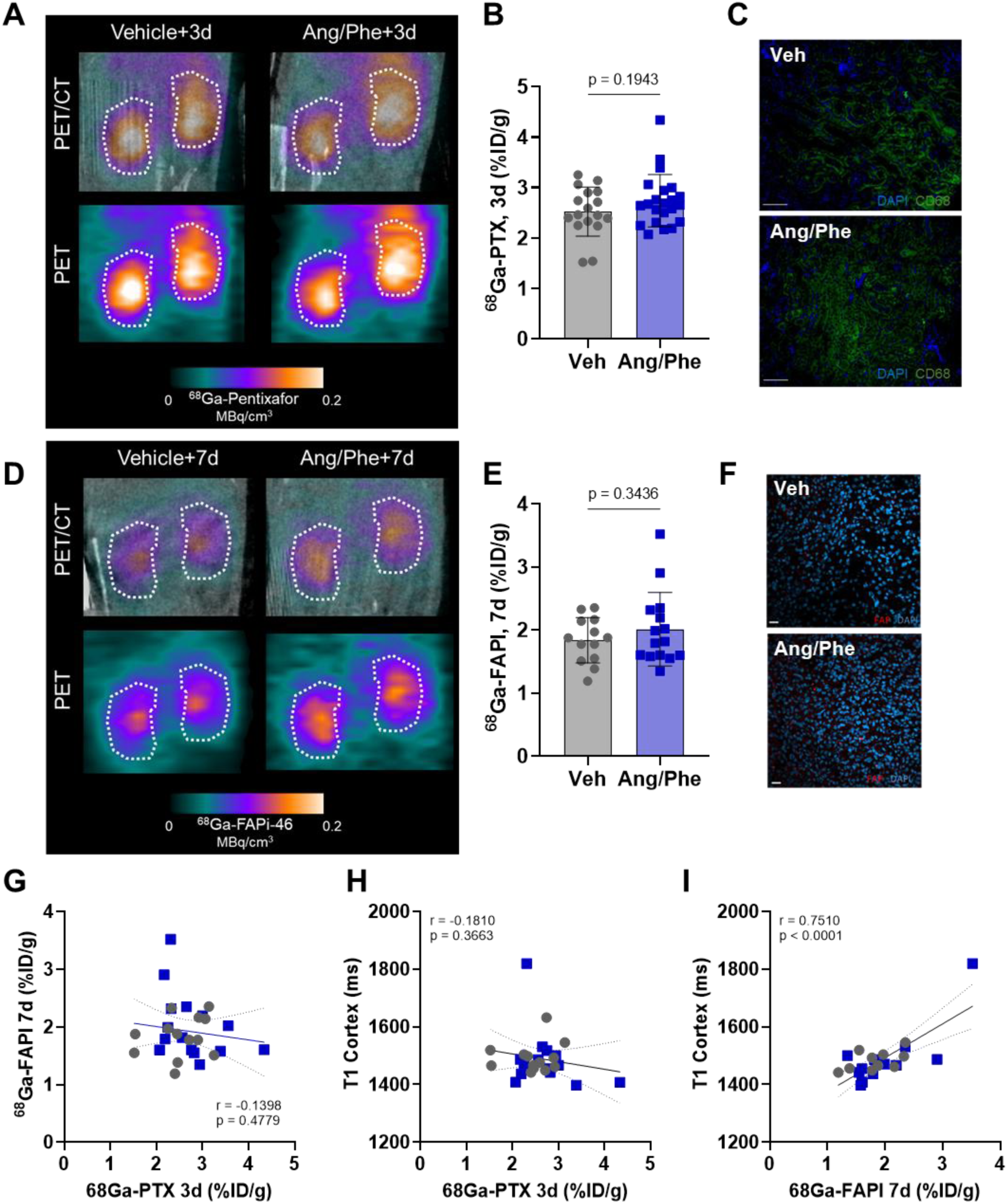
Molecular imaging of inflammation and fibroblast activation in the kidney. (**A**) Representative PET/CT images and (**B**) semi-quantification of ^68^Ga-pentixafor (PTX) signal as % injected dose (ID)/g tissue in the kidney at 3d of angiotensin II/phenylephrine (Ang/Phe) infusion compared to vehicle (Veh) showing chemokine receptor CXCR4 signal (colourscale). (**C**) Immunostaining of CD68 at 3d. (**D**) Representative images and (**E**) semi-quantification of ^68^Ga-FAPI-46 (FAPI) signal in kidney at 7d of Ang/Phe or Veh infusion. (**F**) Immunostaining of fibroblast activation protein (FAP) in renal cortex at 7d. Pearson correlation between (**G**) PTX signal at 3d and FAPI signal at 7d, (**H**) PTX signal at 3d and T1 relaxation at 2 wk and (I) FAPI signal at 7d and T1 relaxation at 2 wk of Ang/Phe or Veh infusion. Statistics: unpaired t test with Welch correction for comparison of two groups, Pearson product-moment correlation between continuous variables.

Imaging with ^68^Ga-FAPI-46 similarly demonstrated a comparable level of FAP signal between Ang/Phe and vehicle infused animals at 7d (Fig 4D,E), which was replicated at 6 weeks (Fig S8). In both Ang/Phe and vehicle groups, the intensity of kidney ^68^Ga-FAPI signal significantly declined from 7d to 6 weeks (Fig S8). Ex vivo autoradiography suggested higher signal in the renal cortex of Ang/Phe infused animals, but was not statistically significant in limited sample size (Fig S6). Consistent with this notion, fluorescence immunostaining revealed detectable FAP in kidneys of Ang/Phe mice at 7d (Fig 4F).

Unlike the heart, correlation analysis did not reveal a clear interrelation between acute inflammation and subsequent fibrosis. The CXCR4 PET signal at 3d did not correlate with the FAP PET signal at 7d (Fig 4G). Likewise, there was no correlation between the CXCR4 signal and prolonged T1 relaxation in the renal cortex at 2 weeks (Fig 4H). However, we observed a significant direct correlation between the FAP signal at 7d and transient prolongation of T1 relaxation time at 2 weeks (Fig 4I), consistent with a predictive value for progressive fibrosis in the kidney after short term Ang/Phe infusion. CXCR4 signal did not predict later reduction in renal perfusion, but the FAP signal directly correlated with renal perfusion at 6 weeks (r= 0.5438, p= 0.0073, Fig S9)

### Immune and fibroblast crosstalk of the heart-kidney axis in hypertensive heart failure

To evaluate inter-organ networking, we next performed correlation analyses between imaging biomarkers and functional readouts in heart and kidney. The CXCR4 signal in the heart directly correlated with the CXCR4 signal in kidney as measured by total body PET imaging (r = 0.4159, p= 0.0084, Fig 5A). This correlation was observed with and without the inclusion of vehicle controls (Fig S8). Conversely, there was no correlation in cardiac and renal FAP signal at 7d (Fig 5B). Since the intensity of cardiac inflammation impacted the severity of fibrosis in the heart, we next evaluated whether the early cardiac CXCR4 signal would correlate with transient kidney fibrosis as measured by T1 relaxation time. Interestingly, the intensity of 68Ga-pentixafor uptake in the left ventricle directly correlated with T1 relaxation time in the renal cortex (Fig 5C), supporting an immune-fibrosis crosstalk across organ systems. Moreover, heightened CXCR4 signal in the kidney at 3d of Ang/Phe infusion likewise predicted cardiac functional decline, with an inverse correlation to left ventricle ejection fraction (Fig 5D).

**Figure 5.**
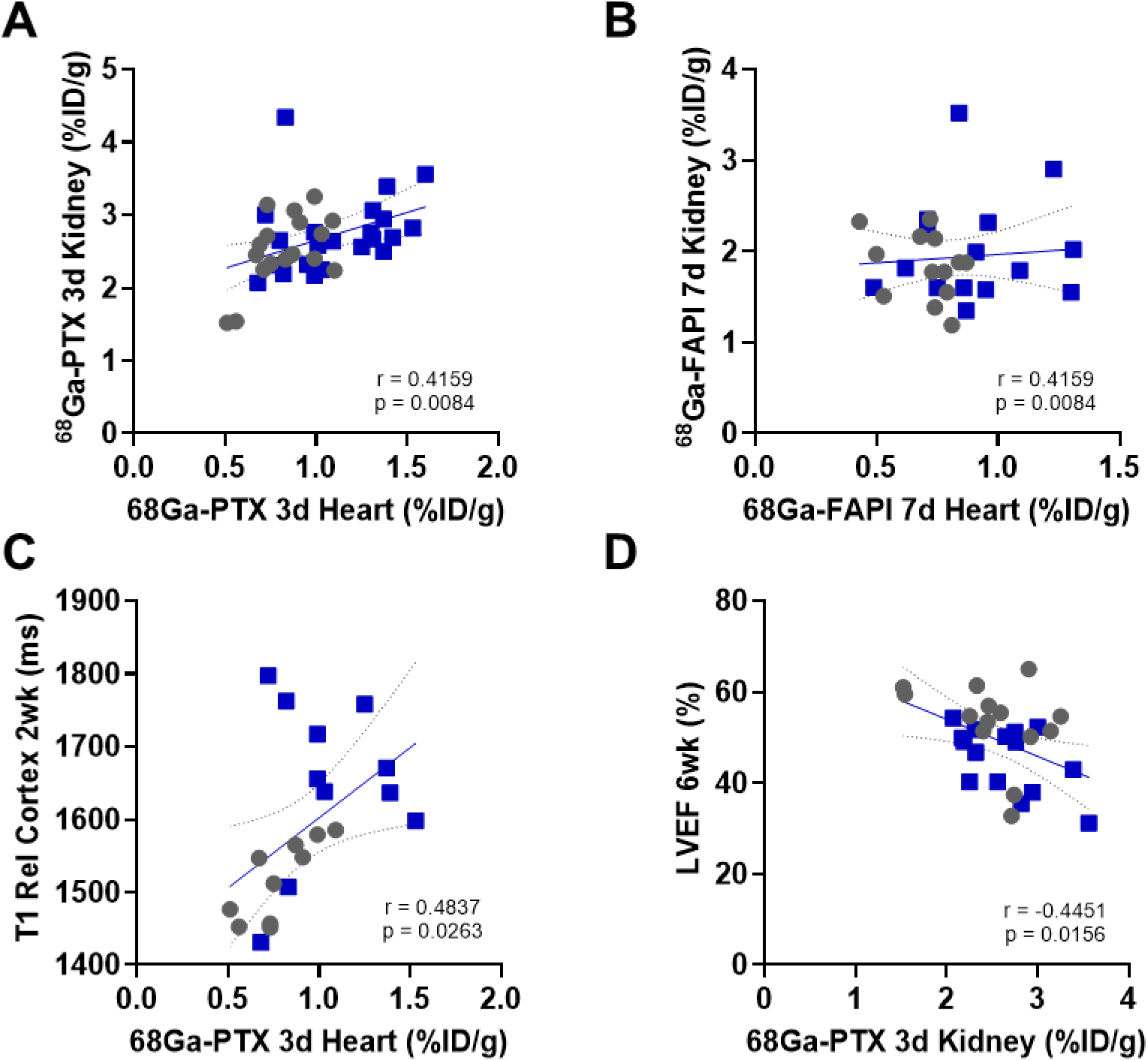
Correlative imaging signal in heart-kidney axis. (**A**) Pearson correlation between ^68^Ga-pentixafor (PTX) signal from chemokine receptor CXCR4 at 3d of angiotensin II/phenylephrine (Ang/Phe) or vehicle (Veh) infusion in heart and kidney. (**B**) Pearson correlation between ^68^Ga-FAPI-46 (FAPI) signal from activated fibroblasts at 7d of Ang/Phe or Veh infusion in heart and kidney. (**C**) Correlation between cardiac PTX signal at 3d and T1 relaxation in renal cortex at 2 wk. (**D**) Correlation between renal PTX signal at 3d and left ventricle ejection fraction (LVEF) at 6 wk.

## DISCUSSION

The immune-fibrosis network has emerged as an intriguing therapeutic target in cardiovascular disease and provides an opportunity for exquisite imaging-based guidance of interventions.^15^ Localized cardiac and systemic inflammation has been proposed as a linking factor in cardiorenal syndrome.(ref) Here, we explored this intersection using total body molecular imaging to characterize both CXCR4 immune cell activity and fibroblast activation in the heart and kidney after temporary hypertension. We observed a persistent decline in cardiac systolic function associated with modest cardiomyocyte hypertrophy and increased left ventricle fibrosis. Imaging of CXCR4 identified diffuse inflammation in the myocardium following 3d of Ang/Phe infusion and subsequent expression of FAP. Conversely, despite subchronic indications of fibrosis in the kidney identified on magnetic resonance imaging, neither CXCR4 nor FAP PET imaging denoted significant changes in response to Ang/Phe infusion. Nonetheless, the intensity of the cardiac inflammation and fibroblast activation signal correlated with signal in the kidney, supporting an immune-fibroblast activation pathway in the progression of hypertensive heart and kidney disease.

Inflammation and fibroblast activation are inherently linked, particularly in the heart.^17^ Following coronary ischemia-reperfusion in mice, there is a close proximity between CD68-expressing macrophages and FAP-positive activated fibroblasts throughout the area at risk of the left ventricle.^18^ In fact, acute myocardial infarction patients with the highest C-reactive protein level tend to have a larger FAP-positive area of the left ventricle early after insult.^19^ Despite more diffuse inflammation in the present study, the heart nevertheless exhibits a similar temporal pattern of immune cell infiltration followed by mobilization of activated fibroblasts. Importantly, FAP has been identified in spatial transcriptomic analyses of clinical dilated cardiomyopathy and chronic heart failure.^11,20^

In a prior study, infusion of Ang/Phe at a comparable dose continuously over 6 weeks generated a surprisingly similar severity of fibrosis in mice, though the progressive decline in left ventricle systolic function was more pronounced. Withdrawal of the Ang/Phe stimulus after 7d resulted in a modest reduction in the left ventricle FAP PET signal at 10d, but did not reverse the extent of fibrosis.^21^ Similarly, in the present study, histopathology-defined fibrosis persisted to the terminal endpoint, supporting the notion that FAP identifies a transient subclass of activated fibroblasts. FAP imaging of the kidney ini chronic hypertension over 4 weeks of Ang/Phe delivery identified a gradual increase in renal signal that was ablated by direct blocking with excess unlabeled precursor or by administration of an angtiontensin receptor blocker.^22^ The maximal temporal pattern of renal Ang/Phe expression was observed at 2 weeks of continuous Ang/Phe delivery, whereas the cardiac signal was maximal at 1 week.^22^ Notably, in the present study, we discontinued Ang/Phe after 7d, which may suggest that sustained FAP expression in the kidney requires the continued stimulus. Nonetheless, the elevation of FAP signal in the heart paralleled earlier reports.^21,22^

Image-based evaluation of the heart-kidney axis has been largely restricted to concurrent evaluation of blood flow. A recent clinical study reported that lower renal blood flow led to substantially higher risk of major adverse cardiovascular events in addition to progressive kidney injury.^8^ However, other studies have reported that coronary flow reserve alone and not estimated glomerular filtration rate could independently predict impaired contractile function.^23^ While these rest-stress perfusion studies highlight the circulatory link between organ systems, they cannot address the molecular underpinnings of cardiac and renal dysfunction.

Radionuclide imaging of the kidney poses a significant challenge, as the renal clearance pathway that is desirable in many radiotracers can obscure specific binding in the target tissue. A recent study established the feasibility of imaging FAP in renal tumours, though the intensity of FAP expression tends to exceed the normal clearance.^24^ Moreover, blocking in Ang/Phe mice have also established specific binding of ^68^Ga-FAPI in the kidney at 50-60min after tracer administration.^22^ Substitution of the radioisotope with a longer halflife positron emitter may enable delayed imaging that reduces background retention from radiotracer clearance.

A surgical mouse model of myocardial infarction demonstrated that the intensity of the CXCR4 PET signal in the infarct territory directly correlated with increased tracer retention in the kidney.^9^ Moreover, severity of contractile dysfunction paralleled higher inflammatory signal in the kidney. Translation to clinical application revealed higher accumulation of 68Ga-pentixafor in the remote myocardium of acute ST-elevation myocardial infarction patients exhibiting worsening renal function.^9^ Short-term anti-inflammatory therapy has been suggested as a pathway to counter worsening cardiac and renal function in chronic heart failure and cardiorenal syndrome.^25^ This concept has not been rigorously tested, though the wider availability of molecular imaging agents targeting immune cells and downstream fibrosis may facilitate patient selection and denote a molecular response to targeted therapy.

The renin-angiotensin-aldosterone system to hypertension in part through modulation of the sympathetic nervous system. In deoxycorticosterone-salt rat model of hypertension, heightened rensal sympathetic nerve activity was associated with rising levels of urinary pro-inflammatory cytokines and macrophage infiltration in the dysinnervated kidney.^26^ Spontaneously hypertensive rats also exhibit rising levels of interleukin-1β, interleukin-6, and tumor necrosis factor-α in the kidney during maturation and development of hypertension.^27^ Rodent models of kidney fibrosis such as the unilateral uretal obstruction and ischemia-reperfusion injury models lead to fibroblast activation and extensive tissue fibrosis.^28,29^ Notably, we report a transient prolongation of renal T1 relaxation time, consistent with kidney fibrosis at the subchronic 2wk timepoint which is subsequently recovered with the removal of the Ang/Phe stimulus. However, magnetic resonance imaging revealed a chronic impairment of renal blood flow which may indicate lasting damage.

Fibroblast activation protein is an attractive therapeutic target given its selective expression in the heart during injury and virtual absence in healthy tissue.^30,31^ Immunomodulatory therapy using chimeric antigen receptor T cells against FAP reversed fibrosis in mice receiving continuous Ang/Phe.^14^ Importantly, the CAR T cells were administered at 1 week of hypertension when interstitial fibrosis was already present, and anti-FAP treatment reversed fibrosis at 4 weeks despite continued Ang/Phe delivery.^14^ In vivo generation of anti-FAP CAR-T cells using a lipid nanoparticle mRNA delivery similarly arrested interstitial fibrosis when delivered at 1 week of Ang/Phe administration, and restored chronic systolic and diastolic function.^32^ More recently, researchers reported on a FAP-targeted vaccine, where three different FAP peptides generated a sustained increased in anti-FAP antibody titre and prevented interstitial fibrosis following 4 weeks of Ang/Phe administration in mice.^33^ Nonetheless, the present study underscores the expression of FAP outside of heart tissue which may generate non-target response to such interventions. Molecular imaging provides a complementary avenue to non-invasively monitor the response to FAP-targeted therapy in non-target organs.

Given the apparent influence of local immune cell activity on FAP, anti-inflammatory approaches may similarly arrest fibroblast activation and tissue fibrosis. In a mouse model of myocardial infarction, short-term treatment with the angiotensin converting enzyme inhibitor enalapril reduced acute cardiac inflammation and lowered the early expression of FAP in the left ventricle.^19^

Some limitations of the present study should be noted. First, experiments were conducted exclusively in male mice, as females display a blunted response to Ang/Phe induced hypertension.^14^ The biology bases of sex-based differences that underlie this susceptibility warrant further investigation, but was not the focus of the present study. Second, we evaluated cardiac function in terms of ejection fraction and geometry but did not assess diastolic measurements. The Ang/Phe model id known to exert increased ventricle stiffness with prolonged E/é ratio and elevated filling pressure.^32^ Despite only a modest decline in ejection fraction, we nonetheless found an association between cardiac signal from both imaging biomarkers with a lower ejection fraction. Third, we did not directly measure renal function using by estimated glomerular filtration rate or creatinine clearance. Prior studies have pointed to the challenge of these measurements in mice,^34^ and the relatively short term hypertension was not expected to generate a robust renal phenotype in the present study. We did identify differences in kidney magnetic resonance imaging parameters, but future studies would need a more expansive workup of kidney function.

In conclusion, hypertension by transient infusion of Ang/Phe leads to inflammation and subsequent fibroblast activation in the heart, and persistent chronic fibrosis despite removal of the hypertensive stimulus. While imaging biomarkers identified these processes in the heart, isolation of renal inflammation and fibroblast activation was hindered by tracer clearance. However, the intensity of cardiac inflammation predicted altered subchronic renal tissue composition and chronic contractile function. Total body molecular imaging may provide valuable insight into the immune-fibrosis network between organs, and exquisitely monitor off-target effects of novel targeted interventions.

## METHODS

### Animals

All experiments were performed in male C57BL/6N mice purchased from Chares River (Sulzfeld, Germany) and housed in groups in a temperature-controlled facility under 14h/10h light/dark cycle with enrichment material and water and standard laboratory diet (Altromin) ad libitum. All animal experiments were approved by local state authorities (Niedersächsisches Landesamt für Verbraucherschutz und Lebensmittelsicherheit).

### Study design

Mice were treated for 7d with Ang/Phe (n=45) or saline (n=32) administered over a transplanted osmotic minipump, as previously described (Aghajanian). Animals were scanned targeting CXCR4 using ^68^Ga-Pentixafor 3d after implantation and targeting FAP using ^68^Ga-FAPI-46 at 7d and 6wks post implantation. MRI was used to assess kidney composition and cardiac function at baseline, 2 and 6weeks after implantation.

### Hypertension model

Ang II (SigmaAldirch/Merck) and Phenylelphrin (SigmaAldrich) were used to create hypertension. Therefore, osmotic Minipumps (Alzet®, model 1007D) were filled with Ang II (1.5µg/g/d) and Phenylephrine (50µg/g/d) diluted in 0.1ml saline. For implantation, adult male C57BL/6N mice (19.4g – 43.3g) were pre-treated with Tramadol (Tramal®, Grünenthal GmbH, Germany) over the drinking water (2.5mg/100ml) beginning the day before surgery and carprofen (Caprosol, cp pharma, 5mg/kg) subcutaneously 20min before surgery. Anesthesia was induced using 3% isoflurane (Isofluran-Piramal, Piramal Critical Care, Germany) in 100% oxygen with a flow of 0.8% l/min. Under anesthesia, mice were transferred to a heated surgery table connected to an inhalation mask for anesthesia supply. After eye protection (5% Dexpanthenol, Bepanthen® Bayer, Germany), shaving, cleaning and disinfecting the back of the animals, a skin cut was performed and a subcutaneous pocket was formed. The pump was inserted with the pumping head towards the mice head. Skin was closed using 6-0 polyethylene terephthalate suture (Polyester-S green, Catgut, Germany) and anesthesia was stopped. Until full recovery mice were kept under infrared and then transferred to their home cage. Analgesia was maintained for five days after surgery via Tramadol (Tramal®, Grünenthal GmbH, Germany) over the drinking water (2.5mg/100ml).

## PET tracer synthesis

### PET Image Acquistion

PET images were acquired using a small animal camera (Siemens Inveon small animal PET, Germany) as previously described ^35^. Anesthesia was induced with 3% isoflurane in 100% oxygen with a flow of 3 l/min. ^68^Ga-Pentixafor (12.7±0.48 MBq) and ^68^Ga-FAPI-46 (12.61±0.53 MBq) were injected as a bolus of 0.1-0.15ml intravenously and flushed by 0.1ml heparinized saline for static image acquisition from 50-60min post injection. Animals were placed prone in either a double or single mouse bed and not moved before end of acquisition. ^18^F-FDG for definition of regions of interest was injected intraperitoneally and a 10min static acquisition was performed 20-30min after injection. Additionally, to PET images, computer tomography (CT) images matching PET images were acquired following PET acquisition immediately without moving mice to allow additional colocalization.

### PET Image analysis

For quantitative PET image analysis Inveon Research Workplace 4.2 (Siemens, Germany) was used. When possible, on all imaging timepoints FDG, gallium and CT images of each animal have been acquired. The FDG image was used to define the regions of interest (ROI) (left ventricle myocardium and kidneys). Once the ROI(s) were defined, they were transferred to the matching ^68^Ga-Pentixafor or ^68^Ga-FAPI-46 image. Exact localization was double checked using CT additionally. Mean voxel intensity per ROI was used to calculate %injected dose per gram (%ID/g) using the following formula: 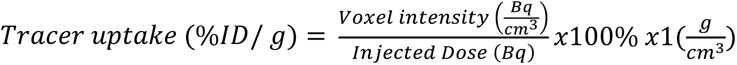

### Small animal MRI

For MRI image acquisition a 7 Tesla MRI system (Pharmascan 70/16, Bruker BioSpin MRI GmBH, Ettlingen, Germany; Software ParaVision 6.0.1) was used. All scans were performed under anesthesia (isoflurane 3.5% induction, 1% maintenance). Mice were positioned prone with the chest on the coil (72mm-diameter volume transmit coil (T11070 89/72 Quad to AD, Bruker BioSpin MRI GmbH) combined with an anatomically shaped four-element mouse cardiac phased-array surface receive coil (T20027V3, Bruker BioSpin MRI GmbH). Respiration rate was monitored regularly during acquisition and kept around 50 breaths per minute. Left ventricle function as measured by left ventricle ejection fraction (LVEF) was calculated using the following formula: LVEF= ((LVV_ED_-LVV_ES_)/LVV_ED_)*100; LVV= left ventricle volume at end diastole (ED) or end systole (ES).

Cardiac function and geometry were evaluated using a navigator-based self-gated 2D spoiled gradient echo sequence (IntraGate FLASH, Bruker Biospin MRI GmbH, Ettlingen, Germany)^36^. For analysis of cardiac function, a short axis cine sequence was obtained (25×25mm field of view, 196×196 pixel matrix size, 0.9mm slice thickness, 0.9mm slice distance, 84.64ms repetition time, 1.84ms echo time, 45° flip angle, 10 reconstructed phases, ∼17 minutes acquisition time) and cardiac function was analyzed using Segment v.40 R12067 (Medviso, Sweden) by manually defining endocardial and epicardial contours to calculate end-systolic and end-diastolic volume, left ventricle ejection fraction and left ventricle mass.

For T1 mapping, a Look-Locker Inversion-Recovery sequence with 2D radial k-space sampling was implemented (35 x 35mm field of view, 128x2048 pixel matrix size, 1mm slice thickness, 3.036ms repetition time, 1.6ms echo time, 1 slice, 15 repetitions, ∼4 minutes acquisition time). According to its cardiac phase (five cardiac phases for the total cardiac cycle) and its inversion time after application of the inversion pulse (64 inversion times, ∼ 7ms – 6600ms) using its direct current signal, data was sorted. In the following, images were reconstructed using a combination of parallel and total variation constraint for the inversion times and the cardiac phases.

For T2 mapping a sequence with the following parameters was applied: 35x35mm2 field of view, 256x256 pixel matrix size, 2mm slice thickness, 1000ms repetition time, 7 echo times (11,22,33,44,55,66,77ms), 1 slice, ∼4min acquisition time.

Arterial spin labeling was used to assess organ perfusion. Therefore, the T1 sequence was used, and additionally the same sequences were measured again but with a non-selective inversion recovery instead of the slice selective inversion recovery. Following, slice selective and non-selective images are reconstructed in the same way as T1 weighted images. Using the difference between both T1 times (T1ss and T1ns), the blood’s T1 time (∼1700ms when scanned with a 7Tesla field strength) and the blood-water partition coefficient (the magnetization ration between arterial blood and perfused time), the blood flow can be quantified using the following formula: 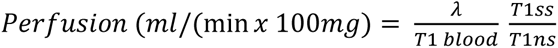

### Ex vivo work up

Organs of interest (kidneys and hearts) were harvested after cervical dislocation. They were snap frozen in cryomolds filled with TissueTek (Sakura Finetek Europe) under liquid nitrogen and stored at -80° until usage. 6 µm thin cryosections were produced using a Cryostat (Leica CM3050 S, Leica Biosystems, Germany) and also stored at -80°C until usage.

### Histology and Immunohistochemistry

To identify collagen in heart and kidneys picrosirius red staining was performed using 500mg picrosirius direct red (Sigma Aldrich) diluted in 500ml picric acid (Sigma Aldrich). For quantification in heart sections, ImageJ (version 1.54g) was used. 4 images per animal in 200x magnification were analyzed and the mean % fibrotic area was calculated. CD68+ cells were identified using a biotin conjugated CD68 antibody (clone MCA1975B, BioRad, Germany).

### Immunofluorescence staining

Tissue from 3d was stained for CD68+ cells using primary CD68 antibody (CD68 Monoclonal Antibody, Invitrogen, #14-0681-82) with secondary antibody (Goat anti rabbit IgG Alexa Fluor 488, Dianova #705-545-003) and DAPI. 7d tissue was stained for FAP using primary FAP antibody (FAP Polyclonal Antibody; Catalog #PA5-51057) and secondary antibody (Donkey anti-rabbit IgG Alexa Fluor 647, Jackson Immuno Research #711-605-152). Cell membrane in cardiac cryosections was stained using wheat germ agglutinin (ThermoFisher Scientific; Catalog #W11261).

### Ex-vivo autoradiography

For ex vivo autoradiography either ^68^Ga-FAPI-46 or ^68^Ga-Pentixafor was injected intravenously. Animals were sacrificed 60min later and organs (heart and kidneys) were harvested and immediately sliced at 15 µm. 30min after excisting the organs, slides were placed onto the film and kept there under room temperature for 60min under light protection. After 60min a gamma counter (CR-35 Bio, Elysia Raytest) acquired autoradiography images to further processing.

### Statistics

All data are presented as mean±standard deviation. Statistical analysis using GraphPad Prism (Ca, USA, Version 10.2.0). For kidney analysis, mean from left and right was calculated and used for further statistical analysis. Two groups were compared using Student’s t-test with Welch’s correction where appropriate. Three groups were compared using one-way analysis with variance (ANOVA) with Šidák’s post-hoc test. Two-way analysis was used for comparing two groups on different time points. To assess the relationship of two specific parameters, Pearson product-moment correlation coefficients.

## Acknowledgement

The authors thank the Preclinical Molecular Imaging, Radiochemistry Laboratories (Nuclear Medicine) and the Small Animal MRI Centre (Central Animal Facility).

