## Supplemental Figures for "Tracking Inflammation and Fibroblast Activation in Hypertensive Heart Failure Across the Cardio-Renal Axis"

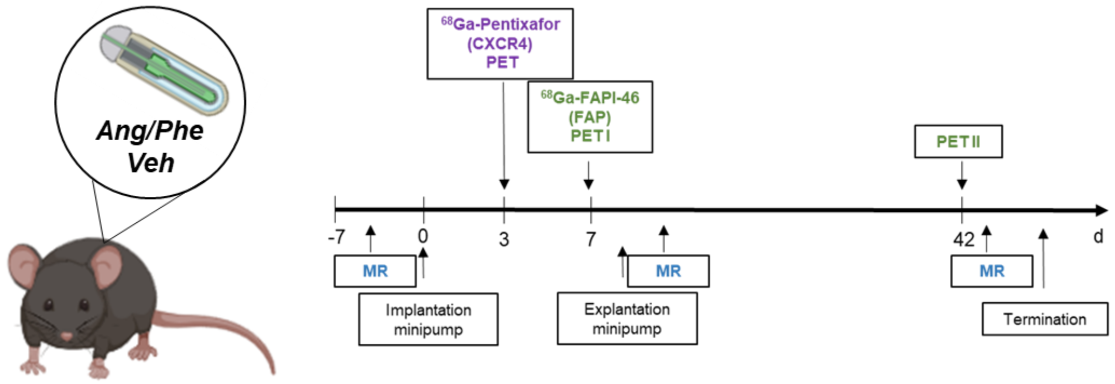


**Suppl. Figure 1. Multimodality and multitracer study design.** Mice are implanted with a subcutaneous osmotic minipump for delivery of angiotensin II / phenylephrine (Ang/Phe) or saline vehicle. Pumps are explanted after 7d of infusion. Magnetic resonance (MR) imaging of heart and kidney is conducted serially at baseline prior to minipump implantation and at 2wk and 6wk after pump implantation and subsequent explantation. Positron emission tomography (PET) images for inflammation targeting chemokine receptor CXCR4 with 68Ga-pentixafor are acquired at 3d after pump implantation. Fibroblast activation protein (FAP) is assessed at 7d and 6wk by $^{68}$Ga-FAPI-46 PET.

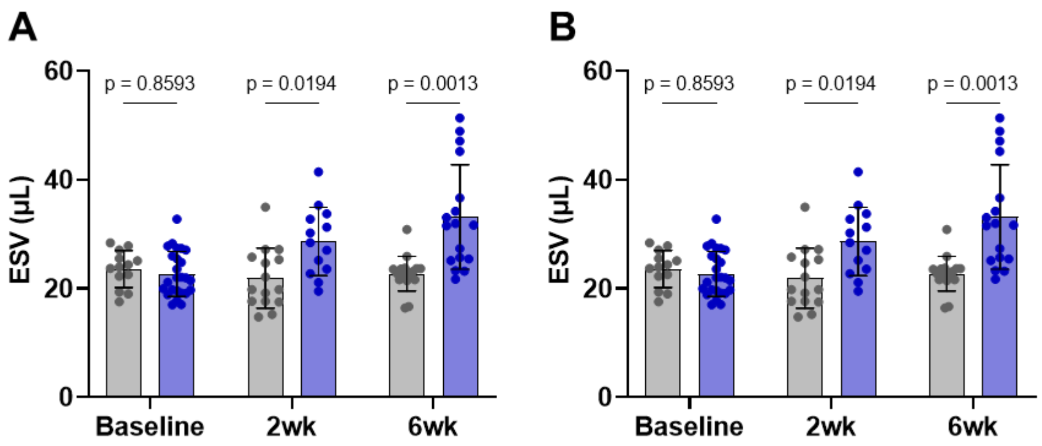


**Suppl. Figure 2. Serial ventricle geometry after 7d angiotensin II/phenylephrine infusion.** (**A**) End systolic volume and (**B**) end diastolic volume in vehicle (grey) and Ang/Phe (blue) delivery from baseline to 6wk. Statistics: mixed effects analysis for multiple comparisons.

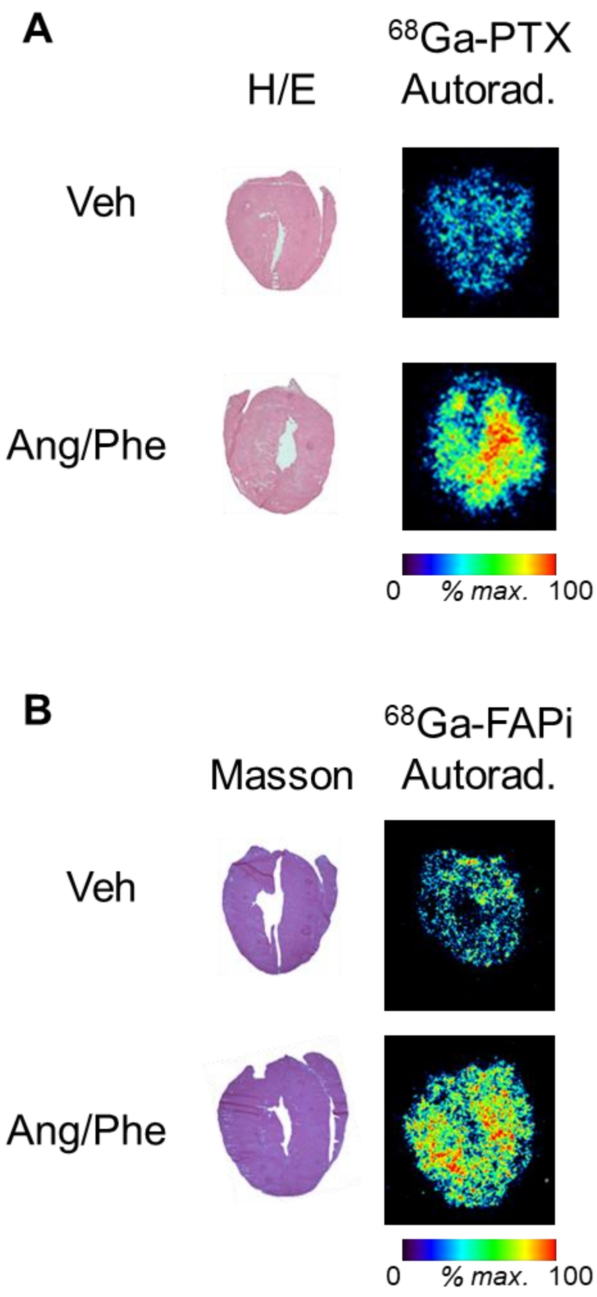


**Suppl. Figure 3. Autoradiography confirms cardiac radiotracer distribution.** (**A**) Masson trichrome staining and *ex vivo* $^{68}$Ga-pentixafor (PTX) autoradiography in adjacent left ventricle sections at 3d of angiotensin II/phenylephrine (Ang/Phe) or saline vehicle (Veh) infusion. (**B**) Masson trichrome staining and *ex vivo* $^{68}$Ga-FAPI-46 (FAPI) autoradiography in adjacent left ventricle sections at 7d of Ang7Phe or Veh infusion.

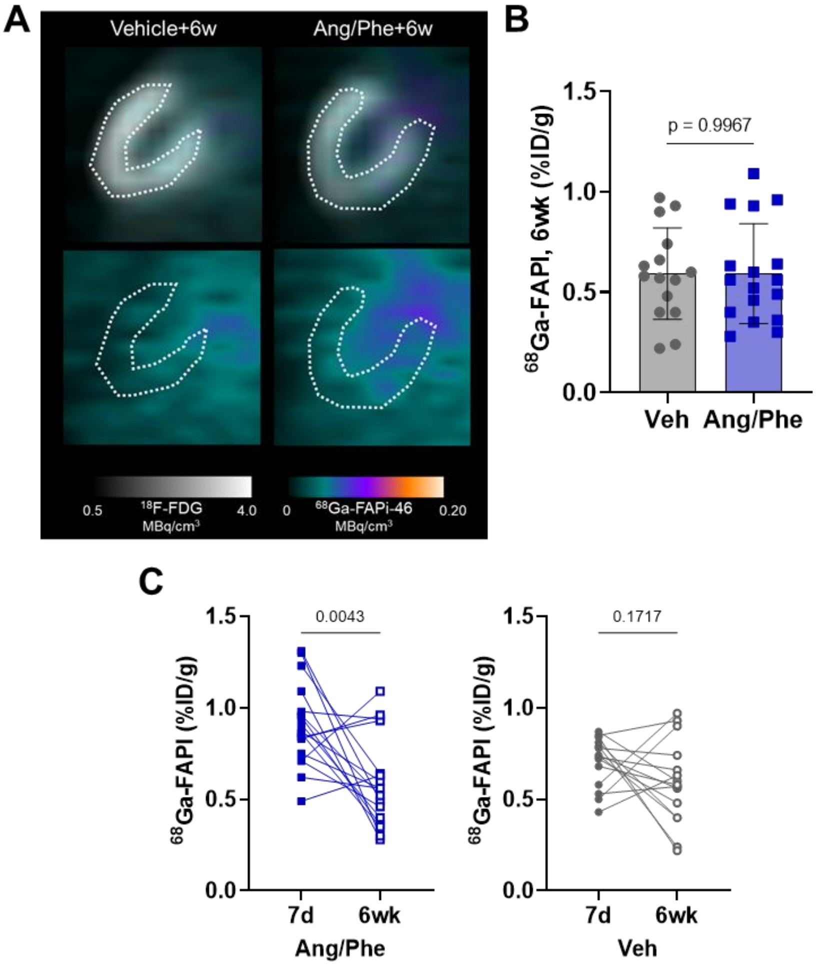


**Suppl. Figure 4. Molecular imaging of chronic fibroblast activation in the heart**. (**A**) Representative images and (**B**) semi-quantification of $^{68}$Ga-FAPI-46 (FAPI) uptake as % injected dose (ID)/g tissue in the left ventricle at 6 wk after transient angiotensin II/phenylephrine (Ang/Phe) infusion compared to vehicle (Veh) showing fibroblast activation protein (FAP, colourscale). Left ventricle contours are defined by subsequent viability imaging with $^{18}$F-fluorodeoxyglcuose (FDG, greyscale). (**C**) Serial comparison of FAP PET signal between 7d and 6 weeks in Ang/Phe (blue) and Veh (grey) mice. Statistics: unpaired t test with Welch correction for comparison of two groups, paired t test for comparison of serial images in individual animals.

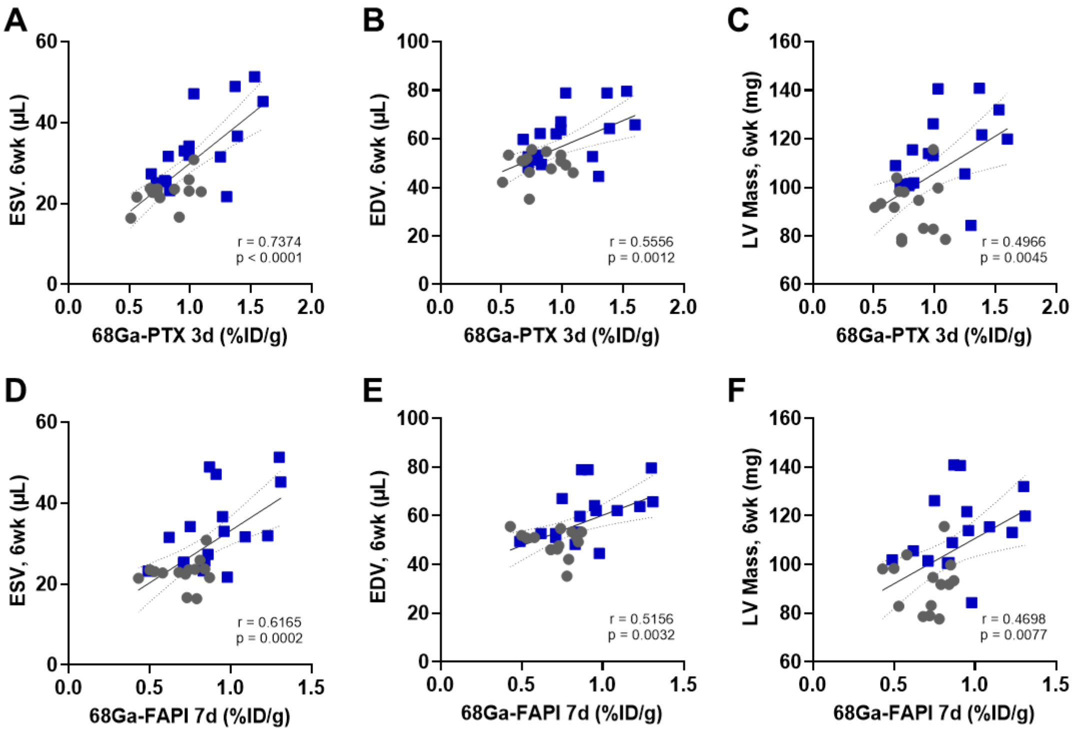


**Suppl. Figure 5. Correlation between cardiac imaging signal and ventricle geometry.** Pearson correlation between cardiac 68Ga-pentixafor (PTX) signal at 3d of angiotensin II/phenylephrine (Ang/Phe) or vehicle (Veh) infusion and (**A**) left ventricle end systolic volume (ESV) at 6 wk, (**B**) left ventricle end diastolic volume (EDV) and (**C**) estimated left ventricle (LV) mass at 6 wk. Pearson correlation between cardiac $^{68}$Ga-FAPI-46 (FAPI) signal at 7d of Ang/Phe or Veh infusion and (**D**) left ventricle ESV at 6 wk, (**E**) left ventricle EDV and (**F**) estimated LV mass at 6 wk.

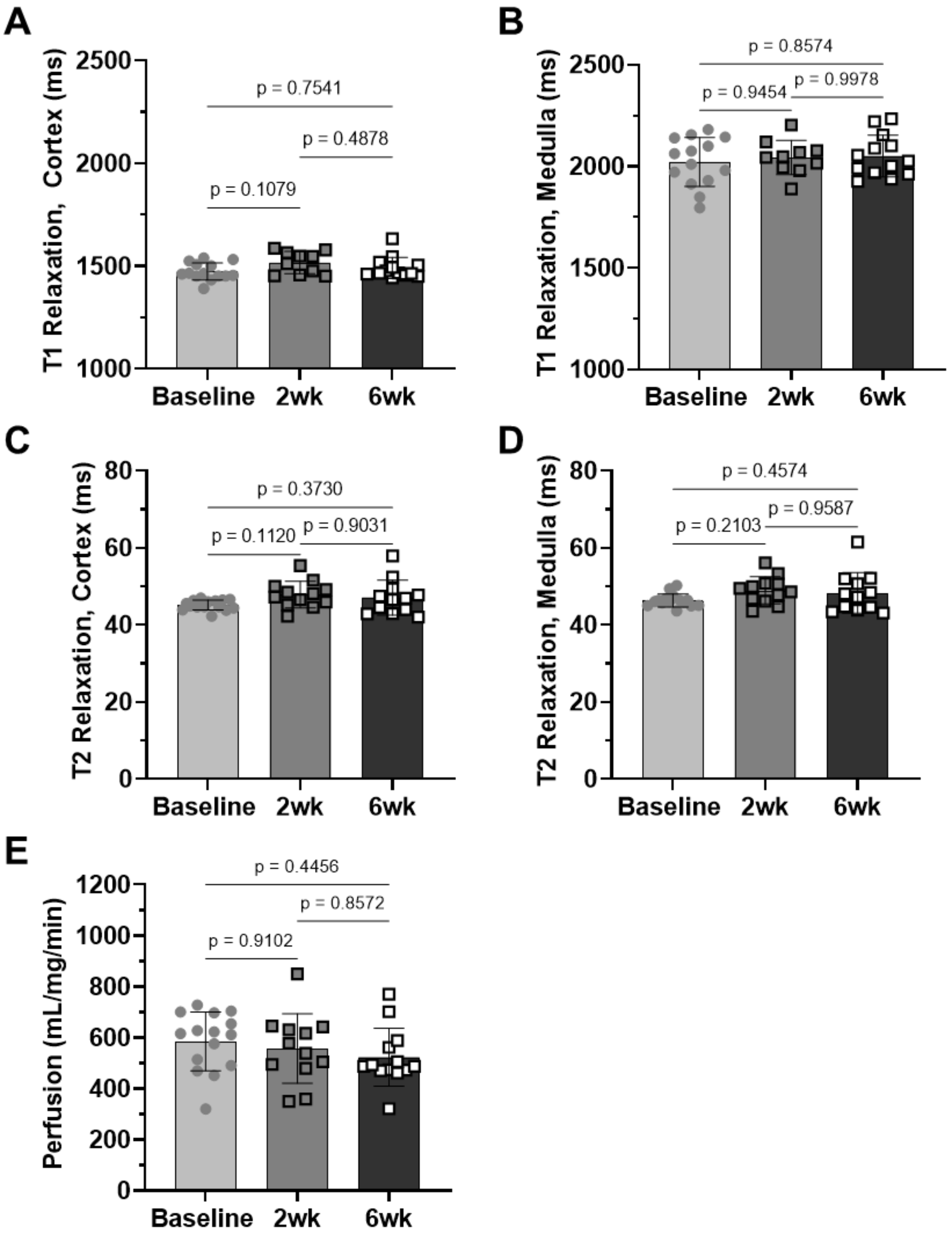


**Suppl. Figure 6. Serial magnetic resonance imaging of kidneys in vehicle infused mice.** Calculated T1 relaxation time in (**A**) renal cortex and (**B**) renal medulla over 6wk after saline vehicle infusion. Calculated T2 relaxation time in (**C**) renal cortex and (**D**) renal medulla over 6wk after saline vehicle infusion. (**E**) Arterial spin labeling derived renal perfusion after Veh treatment. Statistics: ordinary one-way ANOVA for repeated measures.

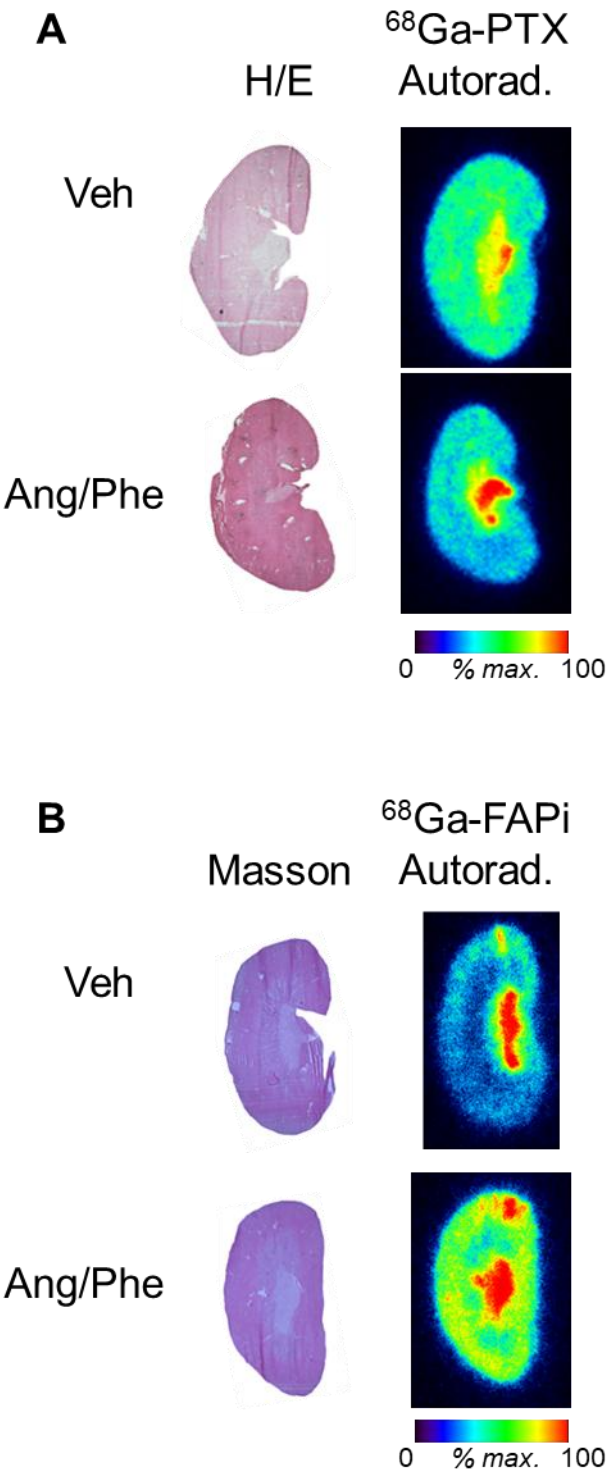


**Suppl. Figure 7. Autoradiography confirms renal radiotracer distribution.** (**A**) Masson trichrome staining and *ex vivo* $^{68}$Ga-pentixafor (PTX) autoradiography in adjacent kidney sections at 3d of angiotensin II/phenylephrine (Ang/Phe) or saline vehicle (Veh) infusion. (**B**) Masson trichrome staining and *ex vivo* $^{68}$Ga-FAPI-46 (FAPI) autoradiography in adjacent kidney sections at 7d of Ang7Phe or Veh infusion.

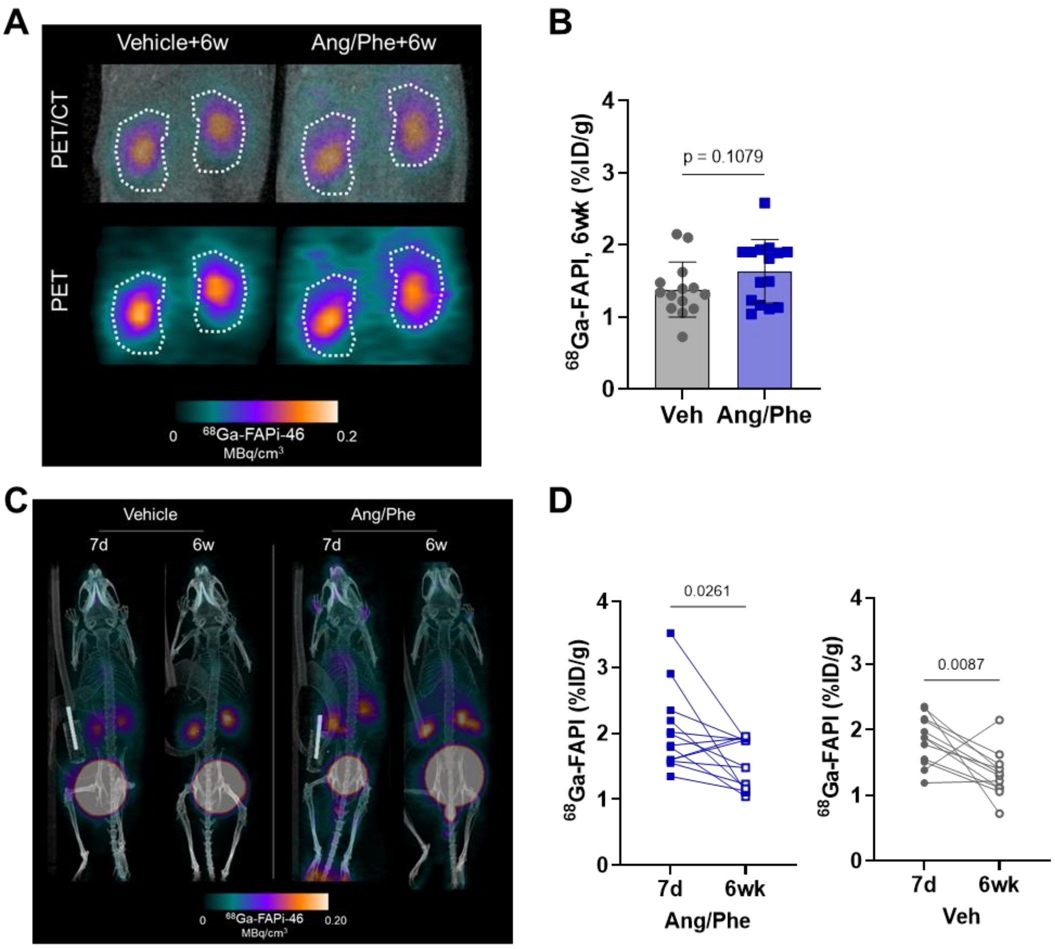


**Suppl. Figure 8. Molecular imaging of chronic fibroblast activation in the kidney**. (**A**) Representative PET/CT images and (**B**) semi-quantification of $^{68}$Ga-FAPI-46 (FAPI) uptake as % injected dose (ID)/g tissue in the kidney at 6 wk after transient angiotensin II/phenylephrine (Ang/Phe) infusion compared to vehicle (Veh) showing fibroblast activation protein (FAP, colourscale). (**C**) Total body maximum intensity projection (MIP) images show intensity of kidney signal over time. (**D**) Serial comparison of FAP PET signal between 7d and 6 weeks in Ang/Phe (blue) and Veh (grey) mice. Statistics: unpaired t test with Welch correction for comparison of two groups, paired t test for comparison of serial images in individual animals.

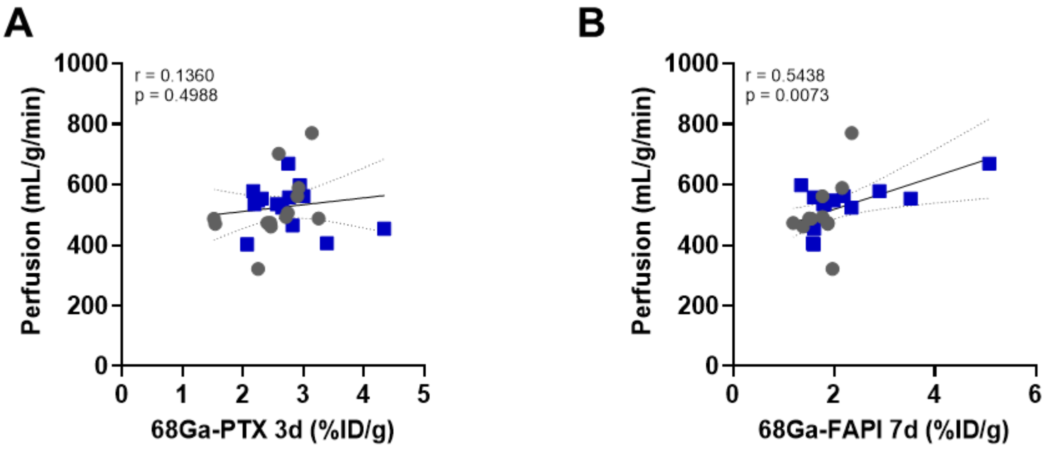


**Suppl. Figure 9. Correlative imaging signal in heart-kidney axis.** (**A**) Pearson correlation between cardiac 68Ga-pentixafor (PTX) signal from chemokine receptor CXCR4 at 3d of angiotensin II/phenylephrine (Ang/Phe) or vehicle (Veh) infusion and renal perfusion at 6 wk. (**D**) Pearson correlation between cardiac 68Ga-FAPI-46 (FAPI) signal from fibroblast activation protein at 7d of Ang/Phe or Veh infusion and renal perfusion at 6wk.
